# Bayesian automatic screening of pneumonia and lung lesions localization from CT scans. A combined method toward a more user-centred and explainable approach

**DOI:** 10.1101/2025.04.08.647710

**Authors:** Álvaro Moure Prado, Alejandro Guerrero-López, Julián D. Arias-Londoño, Juan I. Godino-Llorente

## Abstract

While semantic segmentation allows precise localization of potential lesions, a segmentation based on object detection using bounding boxes is considered more effective for indicating the location of the target without replacing clinical expertise, reducing potential attentional and automation biases.

In this context, this work lays a foundation for better and more explainable localization of lesions in the lungs due to pneumonia by applying a combined approach first using a Bayesian uncertainty measure to evaluate the likelihood of a specific slice being positive in a particular disease and, second, directing the localization of the lesions toward those positive images following object detection methods. This approach provides confidence measures for the model’s predictions and the regions of interest (i.e., bounding boxes) pointing to the lesions, improves the overall model’s explainability, and leaves the expert at the centre of the decision. However, despite its advantages, identifying the existing lesions using bounding boxes also comes with challenges, especially in effectively managing overlapping boxes and developing a fusion strategy to combine smaller boxes into larger ones. In this respect, this paper also proposes a methodology to deal with such a problem.

The methods proposed are tested with a large corpus of approximately 90,000 Computed Tomography (CT) images extracted from public datasets of COVID-19, bacterial, fungal and other viral types of pneumonia, as well as control individuals, paying special attention to the first of these diseases due to the increasing interest raised by the recent pandemic.

For this aim, this work uses a Bayesian neural network based on DenseNet121 for the screening module; and Cascade R-CNN, RetinaNet, and YOLO architectures for identifying the lesions. The method shows screening accuracies for COVID-19 up to 94.61%, which are complemented with robust uncertainty measures. The localization of the associated lesions is carried out with an *mAP*_50_ of 0.5559. The results and the methodology can easily be extrapolated to other pneumonias.

## Introduction

The different types of pneumonia usually leave radiological evidence, which can be observed in those images obtained with Chest X-ray (CXR) and CT, although those obtained with the last technique show clearer traces of the disease, even in its early stages. Thus, CXR and CT have become complementary methods for the screening of pneumonia —including that associated with COVID-19—, not only to improve the specificity but also the sensitivity of the diagnosis (1).

The use of CXR to complement the diagnosis of pneumonia is more extended than CT, mainly due to the availability of basic equipment, the time required for the test, and its cost (2). However, CT imaging offers much higher sensitivity (3) and opens the door to a more accurate localization of lung lesions, which are usually evidenced by lung opacities —typically labelled as Ground-Glass Opacities (GGO)—, consolidations, pleural effusions, and crazy pavement patterns (4, 5) (see Fig. 1).

**Fig. 1.**
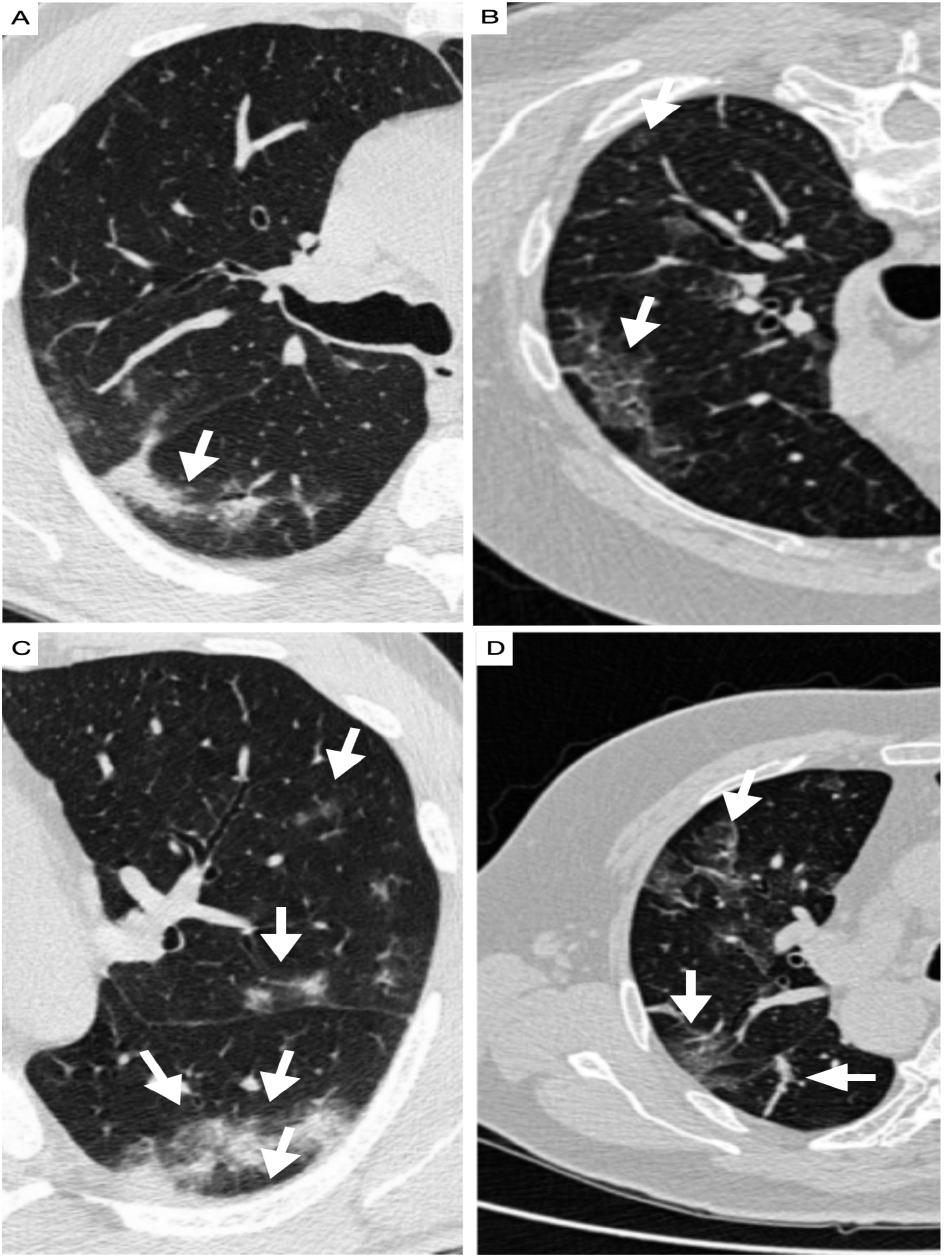
Illustration of the most typical lesions shown in lung CT scans of a COVID-19 patient. GGO are observed in all samples, while C) also shows subpleural effusion and D) consolidations. (Image and annotations obtained from dataset in (6).)

Despite its advantages, analysing every single CT scan is a challenging and time-intensive process requiring specialised expertise. In addition, radiological signs of the different types of pneumonia usually overlap (7), complicating the diagnosis.

Aiming for a simpler and more effective diagnosis process, Artificial Intelligence (AI) assisted frameworks for identifying lung lesions due to pneumonia are valuable tools able to reduce the workload of radiologists while improving the accuracy of the diagnosis and the evaluation of the disease. These systems also have the potential to identify lesions that might otherwise go unnoticed by clinicians.

In this regard, due to the recent pandemic, the academic community quickly recognised this potential not only for the diagnosis of different types of pneumonia but also for the specific case of the pneumonia associated with COVID-19, leading to a significant number of studies combining radiological imaging with various Machine Learning (ML) and Deep Learning (DL) techniques and for identifying the existing lesions in the lungs, primarily using CXR (see (8) and references therein). However, the literature also reports approaches using CT scans. Those works considered most relevant are presented in the following paragraphs, focussing the review on studies that use publicly available datasets, since they are considered more reproducible and are open to a fair comparison (9, 10).

In (11), a lightweight Convolutional Neural Network (CNN) model was employed to differentiate between Healthy Control (HC), non-COVID-19 pneumonia, and COVID-19, achieving an accuracy of 85.03%. The proposed model was trained and tested on a dataset combining samples taken from the COVID-SAN-DIEGO (12) and SIRM (13) databases. The final dataset included 460 CT images (i.e., slices) of COVID-19 cases, 397 of other pneumonias and/or HC subjects. However, it is known that the (12) dataset contains several slices from each patient, but no patient identification is provided. This fact raises concerns about potential biases in the results due to the inclusion of samples from the same patient in the training and testing sets. Another approach in (14) employed a DenseNet for feature extraction and an extreme learning algorithm for classification, obtaining an accuracy of 98.36% also on the COVID-SAN-DIEGO dataset (12). The work in (15) introduced a dataset comprising 9,979 slices negative for COVID-19, 4,001 positive for COVID-19, and 5,705 belonging to HC. A VGG-16 model fine-tuned with a secondary 13-layer network achieved accuracies of up to 92.38% for COVID-19 negative cases. Using the same dataset, (16) trained a custom Wavelet-CNN4 model with a Haar Wavelet-based pooling layer and a Mish activation function, achieving an accuracy of 99.03%. To address irrelevant features in CT scans for COVID-19 detection, the approach proposed in (17) used a DenseNet201 and an Xception as feature extractors, combined with harmony search and adaptive *β*-Hill climbing for feature selection, achieving an accuracy of 97.16%. Another study, (18), made use of an EfficientNet B0 with a voting scheme to exploit correlations between different CT slices from the same scan, achieving an accuracy of 98.99% with samples taken from the COVID-SAO-PAOLO dataset (19) using a 5-fold Cross-Validation (CV). Lastly, (20) compares various CT pre-processing techniques, including Laplacian and Wavelet transforms, adaptive gamma correction, and Contrast Limited Adaptive Histogram Equalization (CLAHE), in conjunction with an EfficientNet, yielding an accuracy of 94.56% using 1,230 images negative for COVID-19, and 1,252 positive. Detailed metrics of these works are summarised in Table 1.

**Table 1.** Overview of the literature in the field for COVID-19 detection and lesion localization using CT. Regardless of the classes chosen in the analysed research, metrics are always averaged by the total number of classes. Data from the studies is not standardized across research papers, therefore the definition of classes may vary from one study to another.

| Ref | Model Architecture | Data | N° sources | Number of cases<sup>1</sup> |  |  | Size | Metrics |
| --- | --- | --- | --- | --- | --- | --- | --- | --- |
|  |  |  |  | HC | Other pneumonia | COVID-19 |  |  |
| (11) | SqueezeNet | (6), (12) | 3 | 397 (slices) | - | 460(slices) | - | $Acc=85.03\%$<br>$Se=87.55\%$<br>$Sp=85.01\%$<br>$F1=86.2$ |
| (14) | DenseNet201 + EL | (12) | 2 | 397 (slices) | - | 349 (slices) | 224x224 | $Acc=98.36\%$<br>$Se=87.55\%$<br>$Sp=85.01\%$<br>$F1=98.25$<br>$AUC=98.36$ |
| (16) | Custom | (15) | 3 | 5705 (slices) | 9979 (slices) | 4001 (slices) | 256x256 | $Acc=99.03\%$<br>$Pr=98.71\%$<br>$Sp=98.01\%$ |
| (15) | VGG16, MLP | (15) | 3 | 5705 (slices) | 9979 (slices) | 4001 (slices) | 256x256 | $Acc=92.2\%$ |
| (17) | DenseNet201, Xception | (19) | 2 | - | 1230 (slices) | 1252(slices) | 224x224 | $Acc=97.16\%$ |
| (18) | EfficientNet B0 (customized) | (19), (12) | 2 | - | 1693 (slices) | 1601 (slices) | 224x224 | $Acc=87.6\%$<br>$Acc=98.99\%$ |
| (20) | EfficientNet B0 | (19) | 2 | - | 1230(slices) | 1252(slices) | 224x224 | $Acc=94.56\%$<br>$Pr=95\%$<br>$Re=91\%$<br>$F1=93\%$ |
| (21) | Mask R-CNN | (22) | 2 | - | - | 750 | 512x512 | $mAP_{50}=0.565$,<br>$mAP_{75}=0.413$,<br>$mAP_{50:95}=0.424$ |
| (23) | Faster R-CNN | (6) | 2 | 1300 (slices) | - | 2000 (slices) | - | $Acc=95.2\%$<br>$Sp=97.4\%$<br>$Se=93.0\%$ |
| (24) | Mask R-CNN | (6), (43) | 2 | - | - | 23860<sup>2</sup> | 512x512 | $DICE=91.97\%$,<br>$Acc=98.39\%$,<br>$Se=97.25\%$ |

<sup>1</sup> Always reported in terms of full CT scans unless otherwise specified.
<sup>2</sup> Expressed in terms of all the individual ROIs inside all CT scans.

Although these studies demonstrate the potential of AI based decision support systems for the screening of different types of pneumonia, including COVID-19, from CT images, these techniques have not yet been transferred to the clinic. This is because their integration into the medical practice requires additional tools supporting the decision-making by pinpointing the regions of the lungs associated with the concomitant lesions. These additional tools would provide a significant degree of explainability to the models by establishing a clear cause-effect relationship.

In this regard, several works have also addressed the localization of lung lesions using two different approaches, which are based on object detection and semantic segmentation techniques.

Initial efforts in this area were focused on object detection approaches using a Bounding Box (BB) to localize the region where each lesion is present. For example, in (21), a Mask R-CNN with a custom classifier module was used to locate and classify lung lesions using a dataset from the China National Center for Bioinformation (22), which includes samples from COVID-19, other types of pneumonia, and HC subjects. The authors reported localization of the lesions with *mAP*_50_ of 0.565, *mAP*_75_ of 0.413, and *mAP*_50:95_ of 0.352% (no Sørensen-Dice coefficient (DICE) values were reported). Furthermore, the architecture concatenated the predicted box coordinates and the confidence scores of the most reliable regions (ranked by confidence) and used them as features for the classification module, achieving an overall precision of 92.22%. The same work also proposed a similar Mask R-CNN model with a custom affinity layer, reporting *mAP*_50_ of 0.614, *mAP*_75_ of 0.414, and *mAP*_50:95_ of 0.424 on the same dataset. In (23), a Faster R-CNN model trained with domain adversarial learning and instance normalisation achieved 95.2% accuracy using the ISMIR (COVID-Seg-1) dataset (6). Furthermore, in (24), a Mask R-CNN with a custom spatiotemporal module achieved a mean DICE score of 91.97 and a mean accuracy of 98.39%. This model used disease stage labels and grouped CT slices into sets of three to improve the localization of the lesion.

More recently, other researchers have addressed the localization of the lesions through semantic segmentation approaches. For example, (25) used a pyramid attention U-Net model on a public COVID-19 dataset (26), achieving a DICE score of 41.28. In (27), a Tomographic LayerCAM model was applied to the Moscow COVID-19 dataset (28), achieving a DICE score of 48. Similarly, (29) used a domain- and content-adaptive convolution on the MIDRC-RICORD-1A dataset (30), achieving a DICE score of 65.10. These works highlight the challenges of pixel-wise segmentation of the lesions on CT scans, which is difficult due to their high variability in shape, size, and location, as well as the difficulty of the manual annotations used as ground truth. This is also a common problem in other application domains such as, for example, pancreatic segmentation (31).

The reviewed studies reporting DICE values show that semantic segmentation approaches not only result in a noticeable decline in performance metrics but are also labour-intensive, as they involve the highly time-consuming task of manually delineating the masks (i.e., the contours of the lesions) used for training. Alternatively, approaching lesion localization using BBs (that is, the rectangular Region of Interest (ROI) including the lesions) requires much simpler annotations. On the other hand, despite being perceived as objective and reliable, decision support systems based on semantic segmentation are prone to generate stronger human inductive biases (especially attentional and automation biases) (32, 33), which can impact final clinical decisions. Thus, decision support systems should only guide the process, leaving clinicians responsible for the delineation of the lesions, overseeing AI outputs, and evaluating potential errors.

In view of the aforementioned and beyond differences in performance and the labelling burden associated with the segmentation masks, there is a significant debate about the relevance and practicality of these two approaches (i.e., object detection and semantic segmentation) in medical practice. Although semantic segmentation provides a precise pixel-level localization of the lesions, the support provided by a localization based on BBs, is considered a relevant factor for the translation of AI based decision support systems to the clinic, since they provide supplementary evidence rather than definitive conclusions (34, 35).

Given the trade-off between annotation effort and precision, the debate on the practicality of object detection versus semantic segmentation in medical practice remains ongoing. While semantic segmentation offers detailed lesion mapping, BBs provide an effective means of directing the clinician’s attention to potential areas of interest without imposing the extensive workload associated with mask annotation. This approach does not seek to replace the expertise of clinicians but rather supports their decision-making process by indicating lesion locations and allowing for more efficient integration of AI-based decision support systems into clinical practice.

This is also confirmed by radiologists, who might prefer advice that acknowledges uncertainty (36) so, tools that communicate their confidence and limitations can prevent over-reliance, especially when physicians are unsure (37). This is also supported by studies indicating that visual annotations, such as marking areas in images, can positively influence diagnostic accuracy (38).

Notwithstanding its advantages, the localization of the lesions using BBs also presents its challenges, particularly in the intelligent handling of overlapping boxes, and in the fusion strategy required to merge small boxes into larger ones.

This is especially important when the lesions are small and scattered throughout the lungs. To date, these challenges are not yet fully solved. Thus, there are no clear guidelines to combine multiple boxes into a larger region, or to leave them as separate ones (i.e., individually bounded areas). Finding the right approach to this issue is crucial to ensure that the system provides meaningful diagnostic support, moving forward with a more explainable decision.

In this regard, the review of the state of the art for pneumonia and COVID-19 screening from CT scans reveals that the explainability issues (39) of the models are scarcely addressed, and are mainly limited to a joint combination of the screening methods with models able to identify the lesions. In general terms, the localization of the lesions is assumed to enhance the explainability of the model by addressing the critical question of “*Why*” a model gives priority to one prediction over another. Furthermore, the level of explainability could be improved by providing reliable confidence measurements about the decisions of the model, allowing users to know “*When they can trust the AI model’s output*” (40). This requirement can be approached by incorporating screening epistemic uncertainty measures to determine if a sample being validated is similar to those seen during training or if there is incomplete information about model parameters. On the other hand, an analysis of potential clustering effects of the screening process is considered crucial to evaluate potential biases in the prediction process that reduce its trustworthiness (e.g., the manufacturer or technology of the scanner, sex, etc.). Furthermore, a differential evaluation of the screening and localization modules is also relevant to improve confidence in automatic decisions. Thus, addressing these aspects is expected to positively impact the perception of the different stakeholders. Based on these insights, the approach presented in this paper focuses on detecting COVID-19 from CT images, followed by the localization of the lesions in those positive cases to improve the explainability of the decisionmaking process. We prioritise the localization of lesions over the segmentation due to its annotation cost-effectiveness and the more relevant role of BBs in assisting radiologists without replacing their expertise. Furthermore, this approach ensures that clinicians are the final decision-makers, reducing potential biases and positioning the system as a tool to enhance their expertise rather than replacing it.

In this respect, the screening phase leverages a Bayesian neural network, which provides not only strong predictive capabilities, but also uncertainty measurements for these predictions (41). In the detection module responsible for locating the lesions in the COVID-19 cases, three distinct DL models are employed: Yolov v8, Cascade R-CNN and RetinaNet. These models had already shown promising results in lung lesion location in previous studies for X-ray images (42). Moreover, a methodology for BB combinations of overlapping and next to each other regions of interest is proposed.

## Supplementary Note 1: Material and Methods

This section describes the material and methods, starting from the datasets, the annotation process to define the ground truth BB labels, the methods for preprocessing the images, the architectures used, and the experimental protocol.

### A. Datasets

To ensure high variability and generalizability, a comprehensive data set was compiled by collecting data from various public sources. In this regard, eight data sets were selected according to the following inclusion/exclusion criteria: (i) the data sets must be publicly available to ensure the reproducibility of the results; (ii) a diverse range of sources (that is, countries) must be represented; (iii) all data sets must include labels indicating COVID-19 status; and (iv) data sets must include annotations indicating the locations of the lesions. Table 2 provides detailed information about the datasets used.

**Table 2.** Summary of the main characteristics of the datasets of CT scans used to train the system. Six out of eight corpora are annotated with the relevant areas affected by the infection. Two total rows are included: one for those datasets used for the *Screening & Localization* (S&L) phase (A–F) and one for *All* (A–H).

| Dataset | Patients | CT Scans | CT Slices | No Lesion / Lesion | Country | Mask | Labels | Experimental Phase |
| --- | --- | --- | --- | --- | --- | --- | --- | --- |
| A (6) | 43 | 100 | 100 | 1 / 99 | Italy | Yes | COVID-19 | Screening & Localization |
| B (6) | 9 | 9 | 829 | 456 / 373 | Global | Yes | COVID-19 | Screening & Localization |
| C (26) | 20 | 20 | 3102 | 1406 / 1696 | Global | Yes | COVID-19 | Screening & Localization |
| D (28) | 1110 | 1110 | 46411 | 45626 / 785 | Russia | Yes | COVID-19 | Screening & Localization |
| E (30) | 102 | 112 | 15887 | 7473 / 8414 | Global | Yes | COVID-19 | Screening & Localization |
| F (22) | 150 | 150 | 21470 | 20921 / 549 | China | Yes | COVID-19 | Screening & Localization |
| G (44) | 169 | 169 | 45471 | 40514 / 4957 | Iran | No | HC, CAP, COVID-19 | Screening |
| H (22) | 1053 | 1400 | 142816 | 107681 / 35135 | China | No | HC, CAP | Screening |
| TOTAL (S&L) | 1434 | 1501 | 87799 | 75883 / 11916 | Global | 6/6 | COVID-19 | Screening & Localization |
| TOTAL (All) | 2656 | 3070 | 276086 | 224078 / 52008 |  | 6/8 | HC, CAP, COVID-19 |  |

#### Dataset A

This dataset comprises 100 CT slices collected by the Italian Society of Medical and Interventional Radiology (SIRM), corresponding to 43 patients. All slices belonging to confirmed COVID-19 cases, were stored in JPG format and resized to 512×512, with some loss of upper-level intensity pixel data. Images were converted to Hounsfield Units (HU) and stored in NIfTI format, resulting in a 512×512×100 array. Annotations were performed using MedSeg^®^ software by an experienced radiologist, marking GGO, consolidations, and pleural effusions as semantic segmentation masks. All slices except one contain annotations for the lesions. The dataset and the processing code are publicly available in (6), though no details about the patient ID, medical history, or recording equipment are provided.

#### Dataset B

This dataset is sourced from (6). It includes 9 volumetric CT scans extracted from Radiopaedia^1^. An experienced radiologist manually identified 373 out of 829 slices as positive for COVID-19. These slices were also semantically annotated using two pixel values, corresponding to GGO and consolidations. This dataset includes slices without infection or annotations. No information about patient characteristics or recording equipment is provided.

#### Dataset C

This dataset, described in (26), comprises 20 full CT scans from the Coronacases Initiative and Radiopaedia. CT slices were manually annotated by junior radiologists (1-5 years of experience) and refined by senior experts (over 5 years of experience) using the ITK-SNAP^®^ package. Annotations included combined right and left lung masks and general infection masks. A total of 3,102 slices were extracted (1,696 with infection masks, and 1,406 with no annotations). The authors reported an average annotation time of 400 ± 15 minutes per scan, typically consisting of 250 slices. No information about patient characteristics or medical equipment was provided.

#### Dataset D

This dataset is part of the MOSMED dataset (28), and includes 1,110 studies from various hospitals in Moscow, covering both positive and negative COVID-19 cases. The studies were categorised into five severity levels based on CT appearances: CT0 (no pneumonia), CT1 (≤ 25% pulmonary involvement), CT2 (25-50% involvement), CT3 (50-75% involvement or rapid progression), and CT4 (≥ 75% involvement with diffuse changes). Fifty scans from categories CT1 to CT4 were labelled with binary infection masks indicating GGO or consolidations. Annotations were performed using MedSeg^®^ software and saved as compressed NIFTI files. Patient distribution is 42% males, 56% females, and 2% unknown, with ages ranging from 18 to 97 years and a median age of 47 years.

#### Dataset E

This dataset is part of the RSNA International COVID-19 Open Radiology Database (RICORD) (30), and includes 120 full lung CT scans from four international hospitals. It was annotated by six thoracic radiologists with an average of six years of experience. The 1a release contains pixel-level labels for infection appearances on all relevant slices, with a section thickness of 2.5-5.0 mm. Slices were reconstructed using a soft-tissue kernel. Image-level annotations include various infectious and noninfectious conditions, while exam-level annotations classify scans as typical, indeterminate, atypical, or negative for pneumonia, among others. Metadata includes the patient’s age, sex, and the number of images per patient. The images were provided in DICOM format, requiring specific windowing for lung tissue visualization. Out of the 120 scans, 112 from 102 patients were usable.

#### Dataset F

This dataset, described in (22), is part of the China Consortium of Chest CT Image Investigation (CC-CCII) and includes CT scans from 4,154 patients, categorised as controls, Community-Acquired Pneumonia (CAP), and COVID-19, totalling 617,775 CT slices. The scans were collected from multiple hospitals across China. After removing errors and noisy slices, 3,993 scans from 2,698 patients remained, comprising 340,190 slices. Additionally, 750 CT slices from 150 COVID-19 patients were manually segmented into background, lung field, GGO, and consolidation by five senior radiologists with 15-25 years of experience. Images were provided in 512x512 resolution as individual JPG files, with segmentation masks available as PNG files. No patient information was provided for the segmented subset.

#### Dataset G

This dataset, described in (44), is the COVID-CT-MD dataset, comprising full CT scans from 76 HC, 60 CAP cases, and 169 positive COVID-19. They were collected by the Babak Imaging Center in Tehran, Iran. Three expert radiologists labelled the scans at the full scan level, individual slice level, and lung lobe level, achieving an overall agreement of 88.9%. Out of all cases, 54 COVID-19 and 25 CAP scans were further analyzed to identify slices with evidence of infection. Annotated findings include GGO, consolidations, crazy paving, bilateral and multifocal lung involvement, and peripheral and lower lung distributions. Patients with these findings and positive Reverse Transcription-Polymerase Chain Reaction (RT-PCR) results were classified as COVID-19 positive. No segmentation masks were available for this dataset.

#### Dataset H

This dataset was extracted from the China National Center for Bioinformation COVID-19 CT Database (22). Class imbalance was addressed by including 700 normal and 700 CAP scans, representing a variety of viral, bacterial, and mycoplasma pneumonia. All scans contain slice-level annotations indicating the presence of infection for CAP cases. This dataset, previously introduced with segmentation masks for COVID-19 samples, does not add new COVID-19 samples but focuses on balancing the representation of normal and CAP classes. This dataset is an extended version of **Dataset F**, and contains only class labels. It is used to balance out the number of HC and CAP in the detection phase.

#### A.1 Data annotation and preprocessing

As previously stated, none of the available datasets contains annotations with the rectangular ROI including the lesions. Therefore, the semantic segmentation annotations available in the compiled datasets (A to F) were converted into BBs. This process was carried out following the next steps: (i) the regions corresponding to lung lesions were identified from the segmentation masks using the connected components algorithm (45) from Scikit-Learn (46). Different lesion types were normalised into a single category to create regions per CT slice; (ii) BB coordinates were extracted from these regions, yielding a list of BBs per CT slice containing lung lesions.

The method faced challenges with multiple small connected components, leading to numerous BBs per slice. To address this, a two-step approach was followed: (i) a variation of the Non-Maximum Suppression (NMS) algorithm was applied to remove completely overlapped boxes, followed by a merging operation for partially overlapped BBs; (ii) Hierarchical Density-Based Spatial Clustering of Applications with Noise (HDBSCAN) (47) was used to find clusters of different densities among the annotations inside the CT. All BBs belonging to the same cluster are then replaced by a single BB which encloses all of them. In HDBSCAN, for any point *x*∈ *D* (where *D* is the dataset) and a given parameter *k* (the minimum samples), the core distance is defined as the smallest radius *ϵ* such that the closed *ϵ*-neighborhood of *x* contains at least *k* points. Formally, one writes

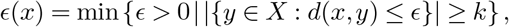

where *d*(*x, y*) is the distance between points *x* and *y*. In the context of BB clustering, a smaller *ϵ* results in fewer BBs being combined, preserving more granular annotations, while a larger *ϵ* results in more BBs being considered part of the same cluster.

Fig. 2 illustrates different examples of CT slices with different pre-processing techniques: NMS only, NMS plus merging, and HDBSCAN with varying parameters. The first two pre-processing methods still provide a relatively large amount of annotations while a larger value of *ϵ* effectively combines further away BBs, resulting in larger annotations which could result in better performance by the object detectors without compromising lesion detection.

**Fig. 2.**
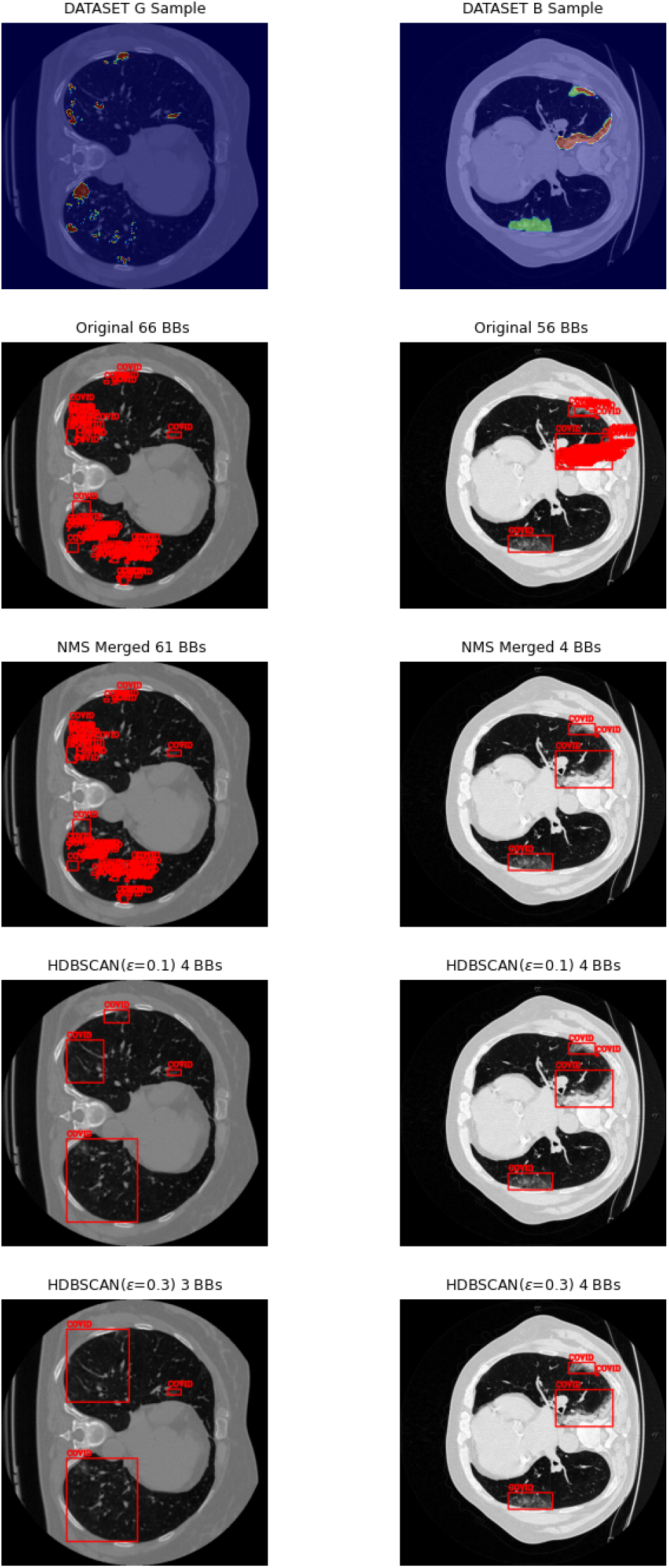
Example images obtained from the final COVID-19 object localization dataset. 4 different pre-processing scenarios are depicted: BB generation without any further pre-process, NMS plus BB merging, HDBSCAN (*ϵ*=0.1), HDBSCAN (*ϵ*=0.3)

This annotation procedure is accessible in the GitHub repository associated with this paper. All images were histogram equalised and CLAHE was used to enhance contrast(41, 48, 49).

#### A.2 Data augmentation

For the detection module, data augmentation is used for samples containing signs of infection to make the model more robust to overfitting. The following augmentation types are used: random rotation, random horizontal flip, random vertical flip, random affine transformation with a translation limit of 10% and scaling limit of 10%, Gaussian blurring with a kernel size of 4 and random sharpness adjustment with a sharpness factor of 2 and random brightness and contrast changes with a limit of 0.1. All enhancements were implemented using the Torchvision^®^ library. Regarding the localization module, mosaic data augmentation is also used for the YoloV8 model, which consists of building patches of multiple training images to generate a mosaic of various training samples.

### B. Analysis Pipeline

Fig. 3 illustrates a simplified pipeline of the overall system. Firstly, the screening module categorises the CT scans according to the health condition (COVID-19, CAP, and HC), filtering out those images belonging to the CAP and HC classes. In a second phase, those categorised as positive for COVID-19 are analysed by the lesion localization module.

**Fig. 3.**
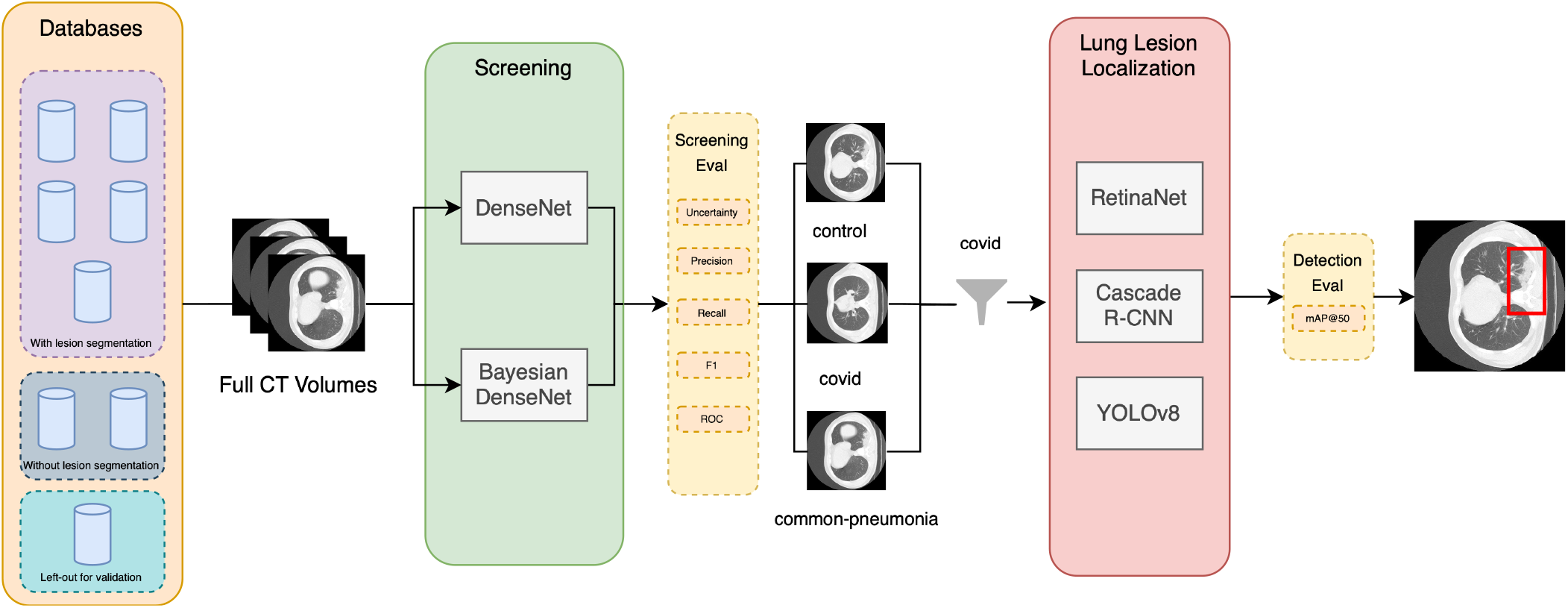
Illustration of the pipeline of the system. A classification module filters out the images corresponding to HC and CAP individuals, while allowing samples categorised as positive for COVID-19 to go through the lesion localization module.

#### B.1 Screening of COVID-19 module

The screening module is based on the approach followed in (41) where a Bayesian adaptation of DenseNet121 was created. Unlike traditional neural networks that use the Maximum Likelihood criterion for regression and classification, Bayesian learning imposes a prior distribution on model parameters and estimates a posterior distribution given the data. This approach enhances the model’s robustness against overfitting and allows it to quantify prediction uncertainty (50).

Training a Bayesian network involves minimising the expected lower bound, which balances two factors: the first measures the difference between the posterior distribution of the network weights and the assumed prior distribution, while the second evaluates the prediction error (50). In this work, the network is implemented using BayesianTorch^®^ (51), which leverages the stochastic gradient descent algorithm to train Bayesian networks through the reparametrisation trick (52), known as “Bayes by Backprop”. During inference, as in (41), the posterior predictive distribution for a given input image is estimated via Monte Carlo (MC) sampling.

#### B.2 Lesion localization module

The lesion localization module leverages three state-of-the-art architectures: You-Only-Look-Once (YOLO) v8 (53), Cascade R-CNN (54), and RetinaNet (55). A key goal of this work is to compare the performance of one-stage detectors (YOLO v8 & RetinaNet) versus a two-stage one (Cascade R-CNN), thereby providing insight into their respective strengths and limitations. These models have been developed for robust performance across diverse datasets and have been widely adopted in the literature for various object detection tasks (42, 56).

All three models are pre-trained using various large datasets such as COCO (57), and then fine-tuned for the task of locating lung lesions in individual slices from CT scans using the previously mentioned datasets.

### C. Evaluation Metrics

The metrics used to evaluate the classification and object detection problems are based on similar principles, though their interpretations differ. The key distinction is that classification metrics assess performance at the image level, while object detection metrics are first calculated at the pixel level and then averaged across images. This difference stems from how “positive instances” are defined for each task. In classification, a positive instance refers to an image belonging to the target class, while a negative instance is any image from a different class (e.g., positive, for COVID-19; and negative, for HC or CAP). However, in object detection, a positive instance is a pixel belonging to a certain structure (e.g., a lung lesion), and a negative instance is a pixel outside it, including the background.

Based on these conventions, True Positives (*TP* ) refer to correctly labelled positive instances, while False Positives (*FP* ) and False Negatives (*FN* ) represent negative and positive instances incorrectly labelled by the system, respectively. Using these definitions, the performance of the CT image classification module was evaluated with the following metrics:

- **Precision (***Pr*) = *TP/*(*TP* + *FP* ), which measures the proportion of true positive instances among the instances classified as positive.
- **Recall (***Re*) = *TP/*(*TP* + *FN* ), which measures the proportion of true positive instances among the actual positive instances.
- *F* 1 **Score** = 2*TP/*(2*T P* + *FP* + *FN* ), which is the harmonic mean of *Pr* and *Re*.
- **Accuracy (***Acc*): measures the proportion of correctly classified instances considering all the classes.

Since the first block in the pipeline addresses a three-class classification problem *Pr, Re*, and *F* 1 score can be calculated for each class and averaged across all three. In this case, *FP* and *FN* refer to instances incorrectly classified, regardless of the specific label assigned. To provide more detailed insights, performance metrics are supplemented with confusion matrices, which display errors broken down by class. Additionally, Receiving Operating Characteristic (ROC) curves and their Area Under the Curve (AUC) are included.

For the object detection system in the second block of the pipeline, evaluation metrics typically involve the DICE coefficient, which is the *F* 1 score at the pixel level, or the intersection over union (Intersection Over Union (IoU)), defined as IoU = *TP/*(*TP* + *FP* + *FN* ). Moreover, since object detection networks assign class labels to each detected BB, these models are often evaluated using mean Average Precision (mAP) at a specific IoU threshold, such as 0.50 (*mAP*_50_).

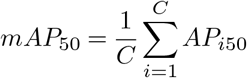

where *C* is the number of classes, and *AP*_*i*50_ is the average precision for class *i* and for the BBs with IoU larger than 0.50.

Additionally, to evaluate the clustering effect of the screening module and to better interpret its ability to separate classes, T-distributed Stochastic Neighbor Embedding (t-SNE) plots for different perplexity values are used to visually represent the feature maps from the last layer (adjacent to the output of the network). Perplexity is considered a crucial hyperparameter, influencing the balance between local and global data structure preservation. These plots are presented with both class and database labels, also helping to identify potential biases related to domain variables associated with the data source (such as hospital, manufacturer of the CT machine, etc.). This approach is inspired by the one presented in (9). Furthermore, the confidence in the prediction of the classification module is evaluated using kernel density estimates of uncertainty measures provided by Bayesian networks, highlighting the differences between correctly and incorrectly classified instances (41) to visually assess the calibration of model uncertainty estimations.

### D. Experimental protocol

This section describes the specific experimental configurations developed to validate each of the two stages of the prediction process: the screening and the COVID-19 lesion localization module.

#### D.1 Screening of COVID-19

Two experimental protocols were developed to validate the viability of the screening module based on a Bayesian model. The first experimental configuration (*Exp1Scr*) consistently labelled all slices from the same CT volume (i.e., patient) according to the health condition, regardless of the presence of lesions. In the second configuration (*Exp2Scr*), only slices showing signs of infection were labelled as CAP, COVID-19, or HC, requiring triple oversampling to balance the dataset. Both experimental configurations were applied to the different architectures tested: the deterministic DenseNet121, and its Bayesian counterpart. All available datasets were used in a Stratified Group 5-fold CV strategy with 80% for training and 20% for validation from each dataset. A group was considered as all slices belonging to the same CT volume for a given patient in a given dataset. This yielded 5 different splits with an average of 1686 *±*11.5 patients for training and 541.2*±* 6.9 patients for validation.

#### D.2 COVID-19 lesion localization

Four experimental configurations were tested to identify the BBs generated by the lung lesion localization module: (i) no BB post-processing at all (*Exp1Loc*), (ii) custom NMS-generated BBs, preserving small instances of the connected component algorithm (*Exp2Loc*); (iii) HDBSCAN BBs with *ϵ* = 0.1, performing unsupervised clustering (*Exp3Loc*); and, (iv) HDBSCAN BBs with *ϵ* = 0.3, where additional BBs were merged into groups (*Exp4Loc*). For all setups, datasets A, B, D, E, and F were used for training. Dataset C was excluded to serve as an independent validation set for the lesion localization module, allowing for an assessment of the robustness of the model on an unseen external cohort. These three experimental configurations were applied to the different object detection architectures tested: Cascade R-CNN, RetinaNet and YOLO v8.

## Supplementary Note 2: Results

This section presents the results obtained. First, for the screening module, and later for the lesion localization one.

### A. Screening of COVID-19

The results of the module for the screening of COVID-19 are summarised in Table 3.

**Table 3.**
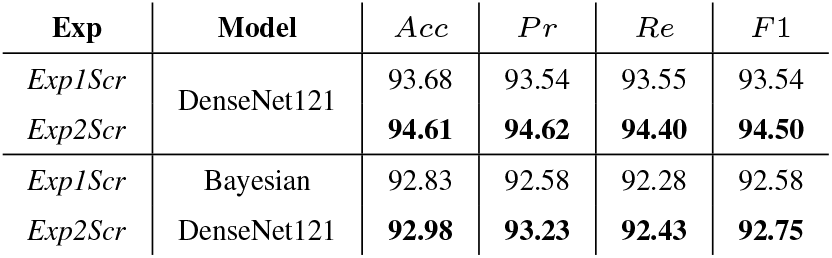
Results of the screening module applying both Bayesian and deterministic networks to the data in *Exp1Scr* and *Exp2Scr*.

In the first experimental configuration (*Exp1Scr*), the deterministic model achieved an *Acc* of 93.68, while the Bayesian counterpart scored 92.83. For comparison purposes, the corresponding confusion matrices and ROC curves are shown in Fig. 4.

**Fig. 4.**
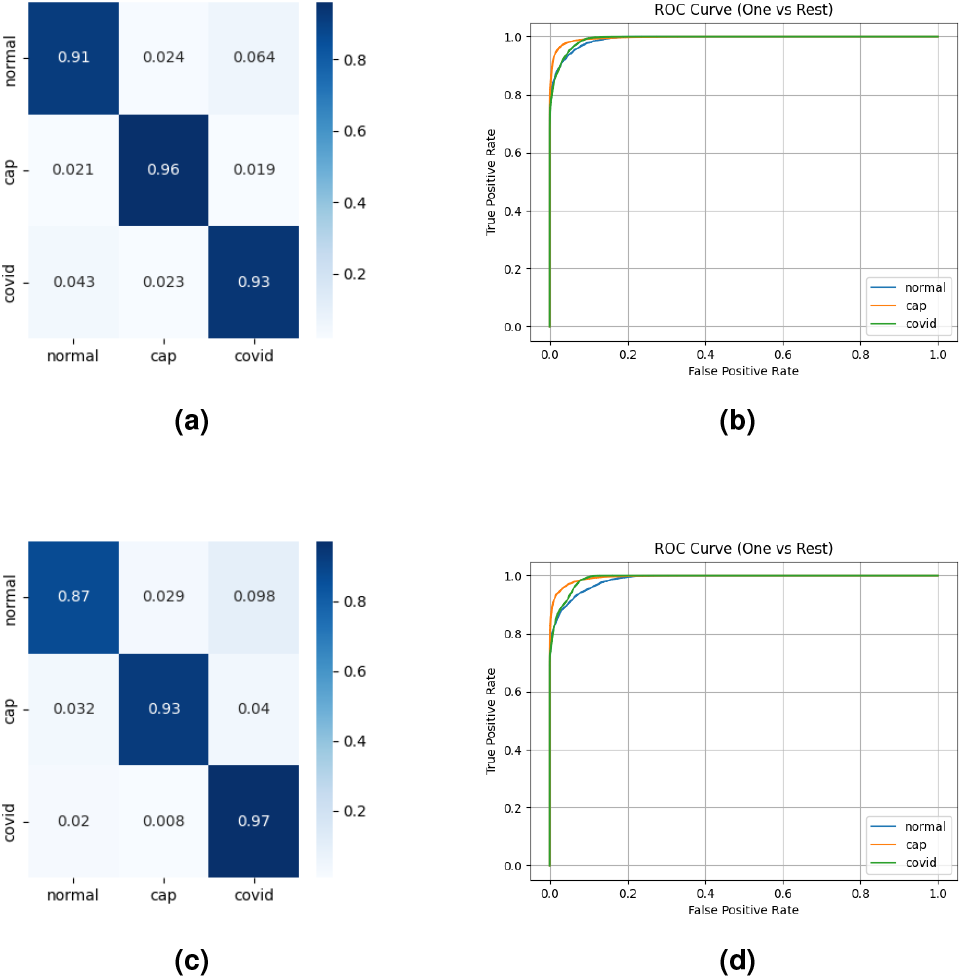
Results for *Exp1Scr* (same labels are assigned to all slices for each scan to a single category): (a) Confusion matrix for DenseNet121; (b) ROC curve for DenseNet121; (c) Confusion matrix for Bayesian DenseNet121; (d) ROC curve for Bayesian DenseNet121.

In the second experimental configuration (*Exp2Scr*), which used only slices with visible infection signs (i.e., with visible lesions) as positive samples, all metrics improved. *Acc* increased 0.93 absolute points, *Pr* 1.08, *Re* 0.85, and *F* 1 score 0.96. The confusion matrices and ROC curves for this experiment are shown in Fig. 5.

**Fig. 5.**
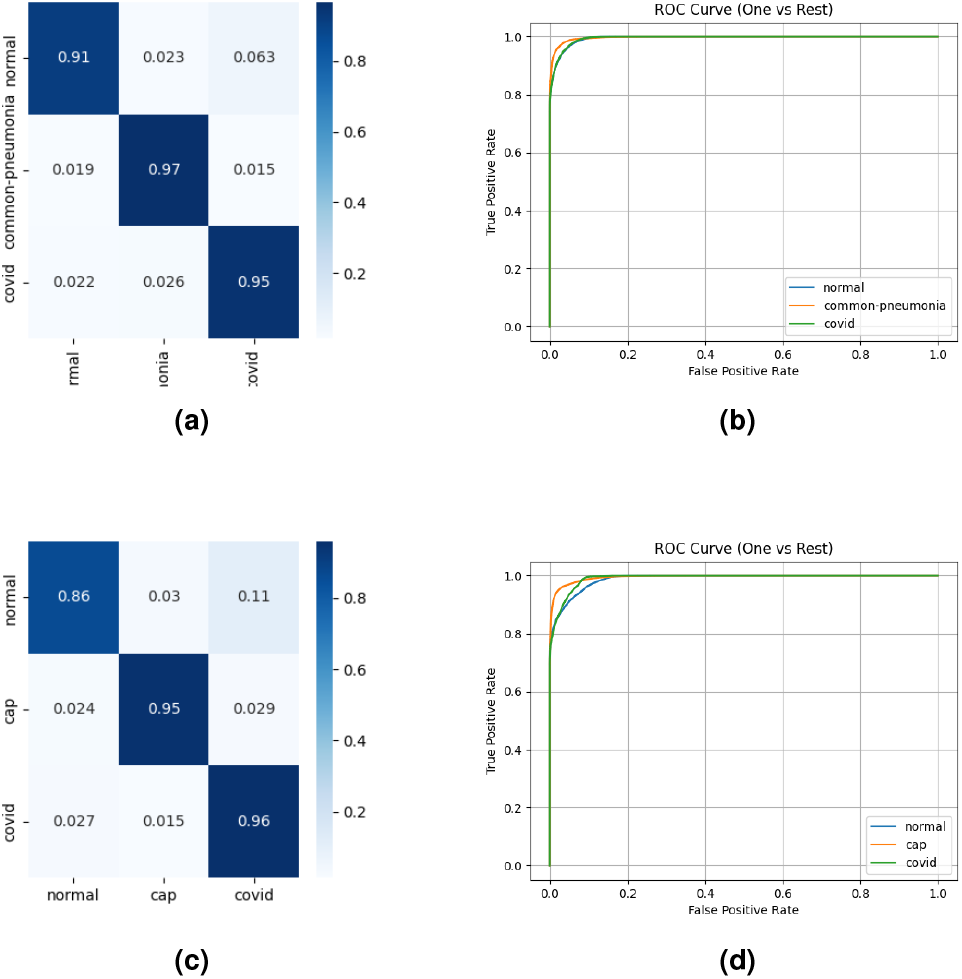
Results for *Exp2Scr* (slices belonging to CAP and COVID-19 classes without infection signs are considered belonging to the HC class): (a) Confusion matrix for DenseNet121; (b) ROC curve for DenseNet121; (c) Confusion matrix for Bayesian DenseNet121; (d) ROC curve for Bayesian DenseNet121.

By comparing the models, the Bayesian version showed a slight decrease in *Acc* of 1.63 absolute points, 1.39 in *Pr*, 1.97 in *Re*, and 1.75 in *F* 1 score. Fig. 6 and 7 compare the predicted uncertainties for both experimental configurations and for the HC, CAP, and COVID-19 classes. Results report no significant changes in uncertainty between the experiments, suggesting potential overfitting to nonspecific characteristics rather than COVID-19-specific ones, as indicated in other studies (41, 58).

**Fig. 6.**
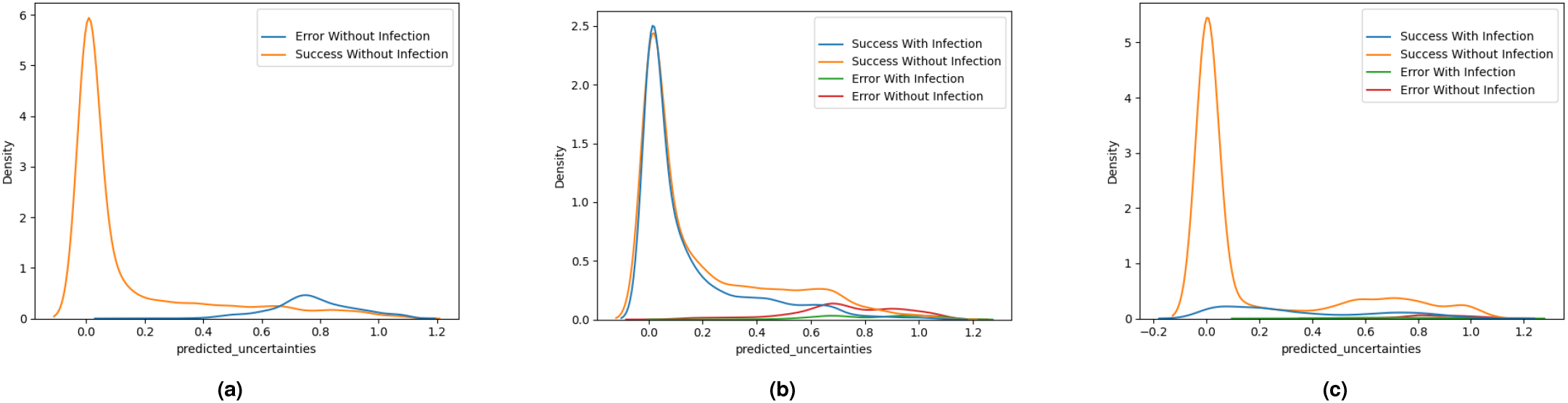
Comparison of predicted uncertainties for the Bayesian version of the DenseNet121 in *Exp1Scr* model for each class. (a) HC; (b) CAP; and, (c) COVID-19. The results are coloured according to whether the sample contained signs of infection labelled by a radiologist or not.

**Fig. 7.**
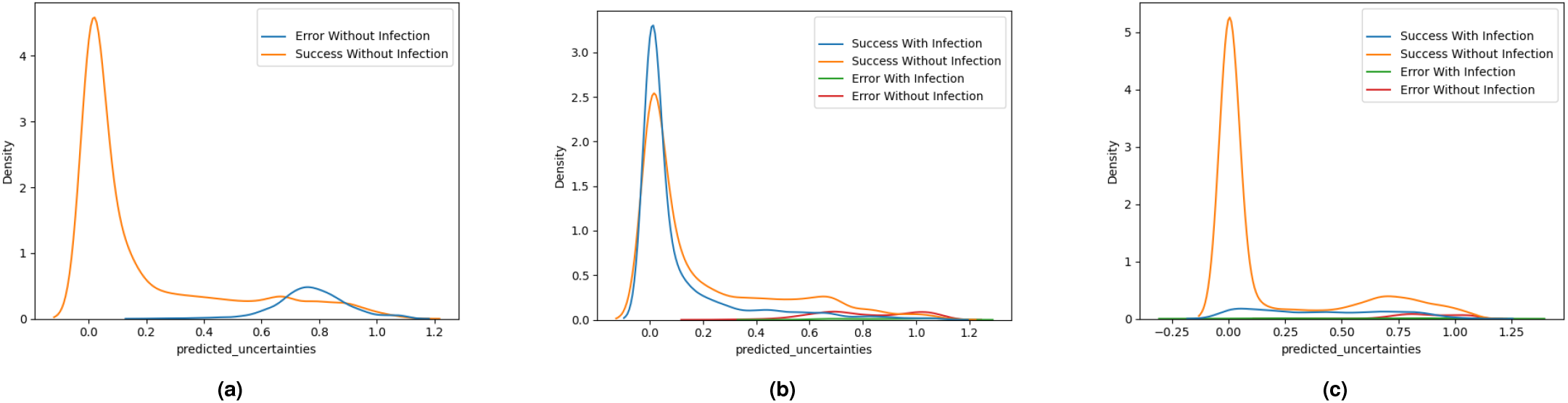
Comparison of predicted uncertainties for the Bayesian version of the DenseNet121 in *Exp2Scr* model for each class. (a) HC; (b) CAP; and, (c) COVID-19. The results are coloured according to whether the sample contained signs of infection labelled by a radiologist or not.

To further assess the performance of the screening model, its potential overfit, and potential biases associated with corpus-dependent characteristics, t-SNE visualisations of the features associated with the last convolutional layer were generated. In Fig. 8a, the projected features are coloured according to the dataset, revealing different clusters, which are similar not matter the perplexity value used to calculate the t-SNE plot. However, in Fig. 8b, where the features are coloured by ground truth labels, class separation is less clear, particularly in datasets such as D (MOSMED) which contains only COVID-19 samples. This suggests that the model might be spatially relying on dataset-specific characteristics rather than disease-specific features, potentially leading to overestimated generalization metrics. Similar problems were identified in (41).

**Fig. 8.**
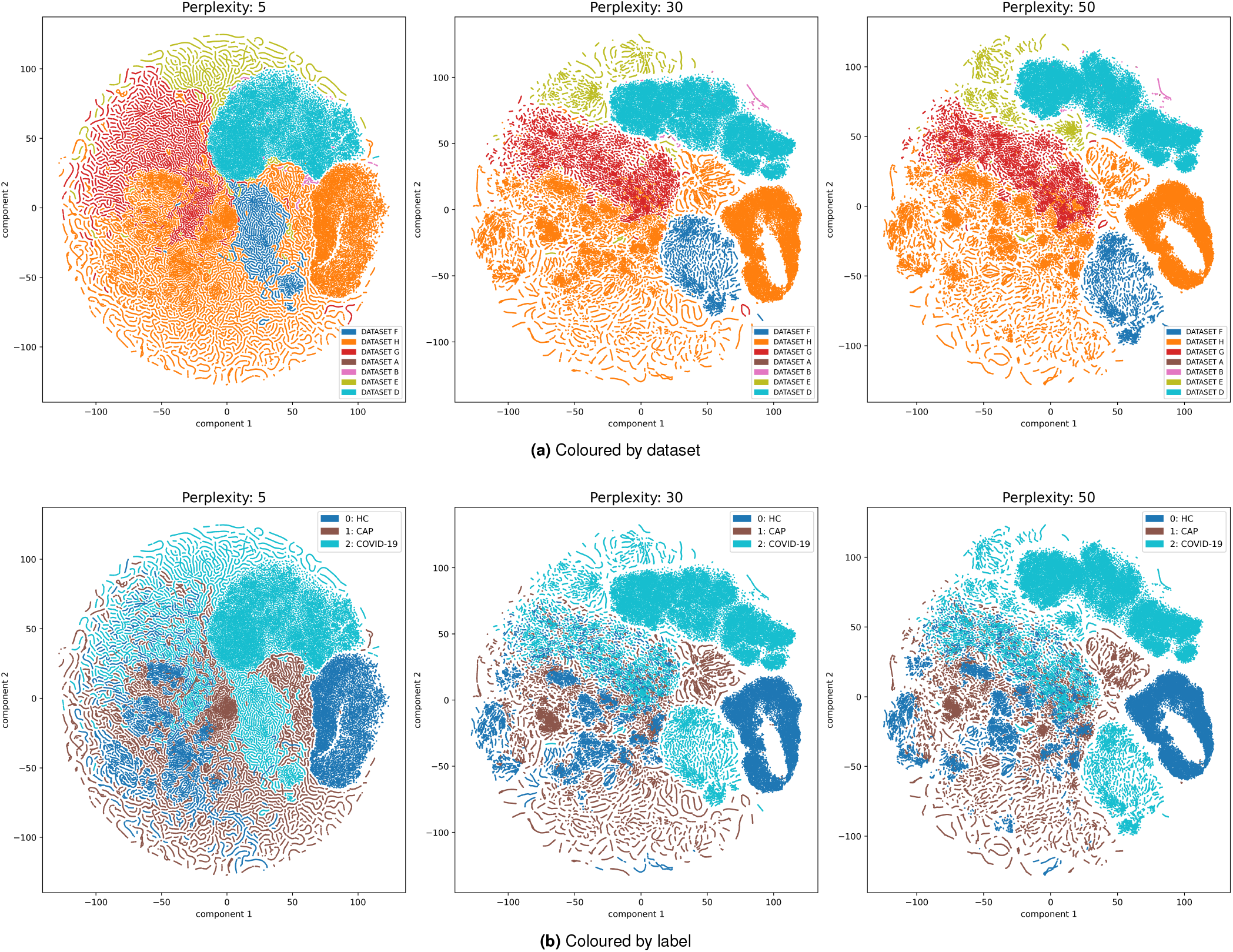
t-SNE representations of the feature maps of the last convolutional layer in DenseNet121 for different values of perplexity. (a) Coloured by dataset; (b) Coloured by label.

The little drops in performance provided by the Bayesian network are widely compensated by the possibility of evaluating the uncertainty of the prediction. Thus, given the results, the Bayesian version of DenseNet121 combined with the *Exp2Scr* experimental configuration has been selected as the final model for the screening module.

### B. Lesion localization

Table 4 summarizes the results of the lesion localization experiments.

**Table 4.** Summary of *mAP*_50_ metrics for the lesion localization experiments for the there models and the different experimental configurations.

| Model | Exp. | BB Pre-Process | $mAP_{50}$ |
| --- | --- | --- | --- |
| Cascade R-CNN | *Exp1Loc* | None | $0.4384 \pm 0.105$ |
| | *Exp2Loc* | NMS Only | $0.5562 \pm 0.061$ |
| | *Exp3Loc* | HDBSCAN ($\epsilon = 0.1$) | $0.5712 \pm 0.056$ |
| | *Exp4Loc* | HDBSCAN ($\epsilon = 0.3$) | **$0.5726 \pm 0.057$** |
| RetinaNet | *Exp1Loc* | None | $0.4028 \pm 0.099$ |
| | *Exp2Loc* | NMS Only | $0.5276 \pm 0.058$ |
| | *Exp3Loc* | HDBSCAN ($\epsilon = 0.1$) | $0.5652 \pm 0.053$ |
| | *Exp4Loc* | HDBSCAN ($\epsilon = 0.3$) | $0.5668 \pm 0.053$ |
| YOLO v8 | *Exp1Loc* | None | $0.4257 \pm 0.042$ |
| | *Exp2Loc* | NMS Only | $0.5253 \pm 0.050$ |
| | *Exp3Loc* | HDBSCAN ($\epsilon = 0.1$) | $0.5591 \pm 0.051$ |
| | *Exp4Loc* | HDBSCAN ($\epsilon = 0.3$) | $0.5580 \pm 0.012$ |

Using HDBSCAN to create larger BBs (*Exp3Loc* and *Exp4Loc*) improved the performance of the localization in terms of *mAP*_50_. This is likely because object detectors typically perform better on medium-to-large instances, while struggling with small objects (59).

Visual inspections of the results further validate these findings. Fig. 9 compares NMS and HDBSCAN (*ϵ* = 0.3) experimental configurations (*Exp2Loc* and *Exp4Loc*), showing that larger BBs generally yield better results (in terms of *mAP*_50_) for larger lesions, although the smaller ones may be missed.

**Fig. 9.**
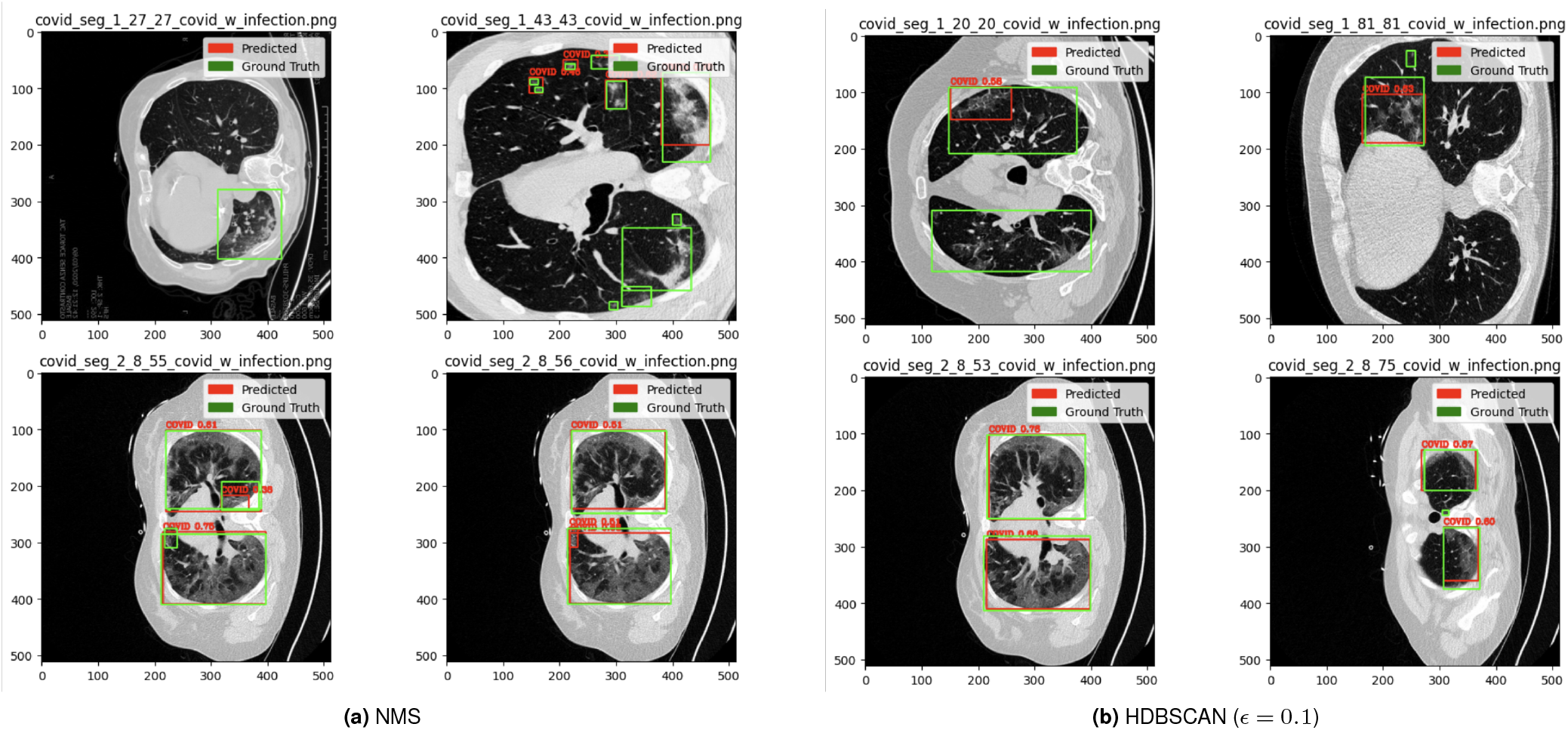
Results of lesion localization using YOLO v8 for: (a) NMS (*Exp1Loc*); and, (b) HDBSCAN (*ϵ* = 0.1) (*Exp2Loc*). Ground truth is in green, and predictions are in red.

Fig. 10 shows a comparison between the NMS and HDB-SCAN pre-processing techniques (*Exp2Loc* and *Exp3Loc*), illustrating the trade-off between detecting smaller versus larger lesions.

**Fig. 10.**
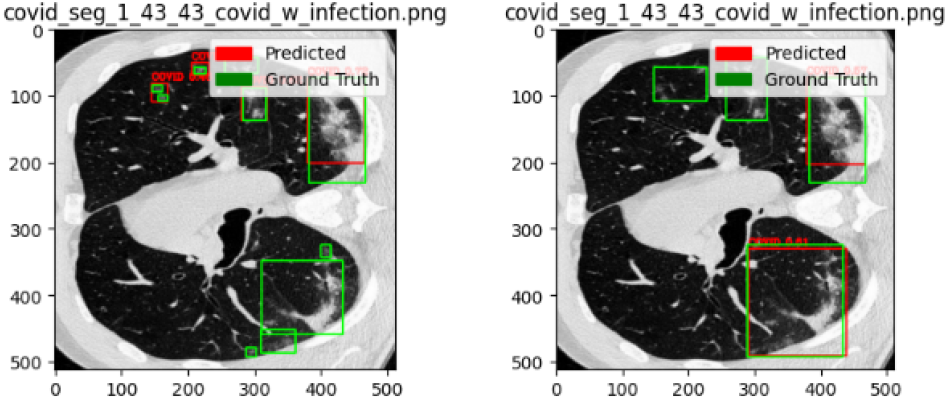
Left to right: Visual comparison of the BBs obtained for NMS (*Exp1Loc*) and HDBSCAN (*ϵ* = 0.1) (*Exp2Loc*). NMS detects smaller lesions, while HDBSCAN captures larger lesions but may miss smaller ones.

In order to validate the results of the lesion localization, an external validation was carried out using dataset C. This additional validation was carried out using the best model previously identified, which is based on a Cascade R-CNN. Results show that the model generalizes well, as shown in Table 5 and Fig. 11, with a slight change for the *ϵ* parameter, suggesting that optimal values are in the range [0.1, 0.3].

**Table 5.**
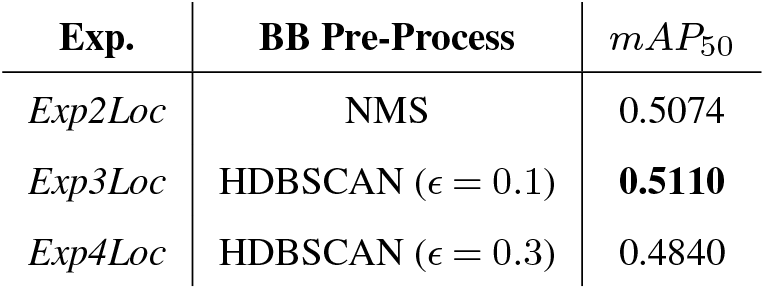
Results on Dataset C using a Cascade R-CNN and the 3 novel proposed BB pre-processing techniques.

| Exp. | BB Pre-Process | $mAP_{50}$ |
| --- | --- | --- |
| *Exp2Loc* | NMS | 0.5074 |
| *Exp3Loc* | HDBSCAN ($\epsilon = 0.1$) | **0.5110** |
| *Exp4Loc* | HDBSCAN ($\epsilon = 0.3$) | 0.4840 |

**Fig. 11.**
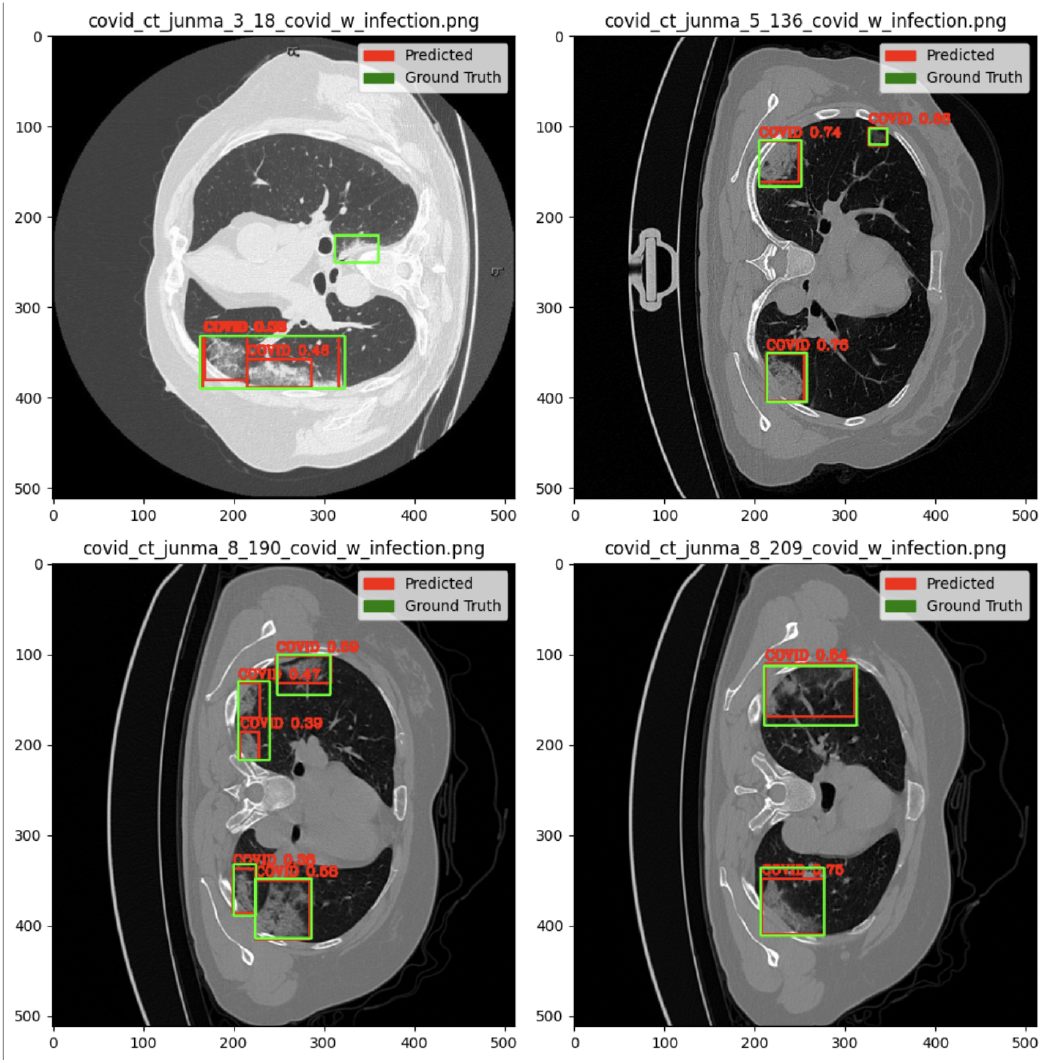
Predicted BBs using the external Dataset C using a Cascade R-CNN and a preprocessing based on HDBSCAN (*ϵ* = 0.1).

## Supplementary Note 3: Discussion & Conclusions

This work presents a combined approach to the design of a reliable screening model for different types of pneumonia and a lung lesion localization system using CT images, addressing the challenges and needs highlighted in the literature regarding explicability and user-centred design. Due to the interest generated by the recent pandemic, special attention was paid to the screening of COVID-19, but results can easily be extrapolated to other pneumonias, as well as to many image-based decision-support systems in medicine.

The approach establishes a solid foundation by first using a Bayesian uncertainty measure to assess the likelihood of a specific CT slice belonging to a certain class, and then focusing on further localization of the lesions for these images. This approach significantly improves the explainability of the screening process, providing confidence measures of the predictions.

The screening was carried out using a three-way classification model, which was developed to distinguish between HC, CAP, and COVID-19. The model was trained with almost 90,000 CT slices extracted from a variety of public datasets, being —to the best of the authors’ knowledge— the largest corpus compiled for this purpose in the state of the art. The classifier was implemented using both a standard DenseNet121 and its Bayesian counterpart, demonstrating strong performance with accuracies up to 94.61% for the best setup. No matter the architecture, training and testing with slices labelled according to the presence (or not) of visible lesions (*Exp2Scr* experimental configuration) improved the classification. On the other hand, the use of the Bayesian approach complements the decisions with uncertainty measures, offering insights into how confident the model is in its predictions, also improving the interpretability of the decision.

While the study acknowledges certain challenges in COVID-19 screening using CT scans, such as validation difficulties and the potential that models rely on spurious features rather than disease-specific characteristics, the overall performance remained robust. This was observed in the t-SNE plots and in the Bayesian model uncertainty analyses, which suggest a certain risk of overfitting to dataset-specific features.

Regarding lesion localization, in order to address the scarcity of publicly available pneumonia (including COVID-19) datasets of CT scans annotated with the corresponding lesions, we employed various pre-processing strategies to generate (and aggregate) the BBs required for training an object detection module. Results report a promising *mAP*_50_ of 0.5559 in lesion localization. Furthermore, when tested on an unseen dataset, the lesion localization module demonstrated consistent performance, closely matching the accuracy reported during internal validation, with only a slight decrease of 0.0449 *mAP*_50_ points. These results underscore the model’s potential applicability in real-world scenarios.

The size of the BBs and the fusing strategy followed have been demonstrated to be a relevant aspect in the design of the system, significantly affecting the accuracy of the lesion segmentation. In general terms, large BBs improve the performance, but a trade-off is required with the usability of the system, since too large boxes provide less relevant information for the diagnosis. To deal with this issue, the method proposed uses a combined approach with NMS, followed by a merging operation for partially overlapped BBs, and HDBSCAN, demonstrating its ability to group BBs into clusters, to combine them into larger boxes, and also to improve the performance of the localization of the lessons. Specifically, results showed the best trade-off between the size of the BBs and the detection of small vs. large lesions for HDBSCAN with *ϵ* = 0.1.

This overall approach supports clinicians by providing not only decisions, but also confidence measures, highlighting the relevant areas of interest in the image instead of providing semantic segmentations (which usually introduce human attentional and automation biases), and without replacing their final decision-making authority, ensuring that the AI tool serves as an aid rather than a substitute in the diagnostic process.

It is also worth noting that the objective of this paper is not to achieve the highest possible accuracy, but rather to propose a framework methodology that emphasizes a more user-centric and explainable approach. While traditional models often prioritize performance metrics, this work shifts the focus toward an explainable and user-centric approach. Ultimately, the goal is to offer a balanced approach that prioritizes both accuracy and the practical, understandable application of the model in real-world scenarios.

However, while these results are promising, further research is needed to enhance its performance and generalizability. Future efforts should investigate semi-supervised learning techniques, such as pseudo-labelling, to refine the training process and further strengthen the model. Additionally, exploring domain alignment and domain adversarial training could be valuable strategies to enhance the model’s generalizability across different datasets and clinical settings (60). While these approaches were beyond the scope of the current study, they represent promising directions for future research.

## ACKNOWLEDGEMENTS

This research was funded by an agreement between Comunidad de Madrid (Consejería de Educación, Universidades, Ciencia y Portavocía) and Universidad Politécnica de Madrid, to finance research actions on SARS-CoV-2 and COVID-19 disease with the REACT-UE resources of the European Regional Development Funds. This work was also supported by the Ministry of Economy and Competitiveness of Spain under grants PID2021-128469OB-I00 and TED2021-131688B-I00, and by Comunidad de Madrid, Spain. Universidad Politécnica de Madrid supports J. D. Arias-Londoño through a María Zambrano UP2021-035 grant funded by European Union-NextGenerationEU. The authors also thank the Madrid ELLIS unit (European Laboratory for Learning & Intelligent Systems) for its indirect support. Finally, the authors gratefully acknowledge Universidad Politécnica de Madrid for providing computing resources on the Magerit Supercomputer.

The source code and trained models are available at the repository: https://github.com/BYO-UPM/CT-COVID

## Footnotes

1 https://radiopaedia.org/

## Notes

https://github.com/BYO-UPM/CT-COVID

### Competing Interest Statement

The authors have declared no competing interest.

